# Preservation of ∼12-hour ultradian rhythms of gene expression of mRNA and protein metabolism in the absence of canonical circadian clock

**DOI:** 10.1101/2023.05.01.538977

**Authors:** Bokai Zhu, Silvia Liu

## Abstract

Besides the ∼24-hour circadian rhythms, ∼12-hour ultradian rhythms of gene expression, metabolism and behaviors exist in animals ranging from crustaceans to mammals. Three major hypotheses were proposed on the origin and mechanisms of regulation of ∼12-hour rhythms, namely that they are not cell-autonomous and controlled by a combination of the circadian clock and environmental cues, that they are regulated by two anti-phase circadian transcriptional factors in a cell-autonomous manner, or that they are established by a cell-autonomous ∼12-hour oscillator. To distinguish among these possibilities, we performed a *post-hoc* analysis of two high temporal resolution transcriptome dataset in animals and cells lacking the canonical circadian clock. In both the liver of BMAL1 knockout mice and *Drosophila* S2 cells, we observed robust and prevalent ∼12-hour rhythms of gene expression enriched in fundamental processes of mRNA and protein metabolism that show large convergence with those identified in wild-type mice liver. Bioinformatics analysis further predicted ELF1 and ATF6B as putative transcription factors regulating the ∼12-hour rhythms of gene expression independently of the circadian clock in both fly and mice. These findings provide additional evidence to support the existence of an evolutionarily conserved 12-hour oscillator that controls ∼12-hour rhythms of gene expression of protein and mRNA metabolism in multiple species.

## Introduction

Biological rhythms are crucial for life on earth and have provided evolutionary advantages to all living organisms, from bacteria to mammals including humans. These rhythms encompass diverse oscillators that regulate different aspects of biology, including the cell cycle that controls cell-division, infradian (with period longer than 24 hours) hormone cycles that regulate the timing of organismal development, and circadian and ultradian (with period less than 24 hours) oscillators that regulate organismal activity, physiology, metabolism, and cellular activity (1-3).

The circadian clock regulates daily cycles in many organisms and is based on a molecular transcriptional-translational feedback loop (TTFL) involving core circadian clock genes of BMAL1, CLOCK, PER and CRY in mammals (4). The TTFL is responsible for establishing and maintaining cell-autonomous 24-hour rhythms of cellular gene expression, metabolic processes, and organismal behavior (4-6). In addition to 24-hour circadian rhythms, biological rhythms have also been shown to have ‘harmonics’ -rhythms with frequencies that are positive integer multiplier of 1.15*10e-5 Hz (24 hour period), such as those that cycle with periods close to 12-hour, 8-hour or 4-hour (7, 8). Oscillations cycling at harmonics frequencies in mammals were first comprehensively characterized in mouse liver using high-resolution temporal microarray in 2009 by Hughes and colleagues (9). Analysis of these data with both Fisher’s G and COSOPT methods yielded a few hundred genes cycling with an approximately 12-hour period (9). Since then, several studies have validated this initial discovery and further expanded the repertoire of ∼12-hour mouse hepatic transcriptome to thousands of genes (10, 11). Strongly contributing to this expansion is the development of novel analytical methods capable of unmasking ultradian from superimposed/co-expressed circadian oscillations (10, 12-14). In addition to liver, wide prevalence of ∼12-hour rhythms of gene expression were further identified in murine white adipose tissue, brown adipose tissue, adrenal gland, aorta, muscle and cornea using multiple mathematical/statistical methods (7, 10-12, 15-25). Using spectrum analytic methods including Fourier, wavelet, autocorrelation analysis and eigenvalue/pencil, ∼12-hour rhythms of locomotor activity, energy homeostasis and/or feeding rhythms were also uncovered from rodents (8, 11, 26).

Gene Ontology (GO) analysis of ∼12-hour transcriptome in different mouse tissues revealed strong enrichment in the entire central dogma information flow (CEDIF) process, ranging from transcription initiation, mRNA processing and export, ribosome biogenesis, translation initiation to protein folding, degradation, processing and sorting in the endoplasmic reticulum (ER) and Golgi and include both anabolic and catabolic processes (15-17). More importantly, in a most recent endeavor, using time of death information as a surrogate for circadian time, ∼12-hour rhythms of gene expression were identified in the dorsolateral prefrontal cortex regions of normal human subjects (27). Intriguingly, human subjects with schizophrenia lose ∼12-hour rhythms in genes associated with unfolded protein response (UPR) and neuronal structural maintenance in the same brain regions (27).

While the evidence supporting the existence of ∼12-hour rhythms in mammals is compelling, the origin and mechanisms of regulation of these rhythms are poorly understood and have been the subject of several theories over the past decade. One of the early hypotheses suggested that these ∼12-hour rhythms are not cell-autonomous and instead established by the combined effects of circadian clock and fasting-feeding cues (9, 28). This hypothesis was mainly supported by the observation that several ∼12-hour rhythms of proteostasis gene expression were disrupted in *Cry1/Cry2* double knockout mice and/or during altered feeding regimen in wild-type mice (28). Alternatively, it was suggested that two circadian transcription factors with anti-phasic transcriptional activity and binding cooperativity are theoretically capable of establishing 12-hour rhythms of gene expression in a cell-autonomous manner (29). However, our recent work challenged these two paradigms, and instead proposed the existence of a dedicated and cell-autonomous 12-hour oscillator responsible for the establishment and maintenance of a large proportion of ∼12-hour ultradian rhythms (11, 15, 16, 20, 21). At the center of the 12-hour oscillator is spliced form of XBP1 (XBP1s), a transcription factor (TF) previously characterized as one of the master regulators for UPR. In XBP1 liver-specific knockout (XBP1 LKO) mice, ∼85% of the ∼12-hour hepatic transcriptome was either abolished (54%) or dampened (31%) compared to wild-type mice, while the core circadian clock genes and most circadian clock-controlled output genes (66%) remained robustly oscillatory with a ∼24-hour period (16). Further, cell-autonomous ∼12-hour rhythms of *Xbp1s* and several representative proteostasis gene expression were observed in synchronized mouse embryonic fibroblasts (MEFs) with or without *Bmal1* knocking down (11, 16).

While these findings support the existence of an XBP1s-dependent 12-hour-oscillator, it remains largely undetermined to what extent these ∼12-hour rhythms of gene expression are preserved in the absence of circadian clock *in vivo* and *in vitro*. In our prior study, while we did observe that many of these ∼12-hour rhythms of gene expression are visibly intact in the liver of BMAL1 knockout (30) and CLOCK ^Δ19^ mutant (31) mice (11), the short duration (24 hours) of these two temporal transcriptome datasets prevented us from rigorous and unbiased identification of all ∼12-hour transcriptome resistant to circadian clock ablation. Herein, utilizing a recently published high temporal resolution/duration (4-hour interval for a total of 48 hours with quadruplicates at each time point) hepatic RNA-seq dataset from whole body BMAL1 knockout mice kept under constant darkness (32), we uncovered widespread hepatic ∼12-hour rhythms of gene expression implicated in mRNA and protein metabolism. We further corroborated our findings with an additional cellular model, the *Drosophila* S2 cells which do not express canonical circadian clock genes (33), and identified hundreds of ∼12-hour transcripts involved in protein and mRNA metabolic pathways. Our findings thus provide further support for the existence of an evolutionarily conserved 12-hour oscillator regulating ∼12-hour rhythms of genetic information flow.

## Result

### ∼12-hour ultradian rhythms of gene expression are prevalent in the liver of free-running BMAL1 knockout mice

To investigate the extent to which ∼12-hour rhythms of gene expression are present and functionally preserved in the absence of the circadian clock *in vivo*, we performed a *post-hoc* analysis of a recently published temporal RNA-seq data collected from the liver of male whole body BMAL1 knockout mice (3 months of age) (34) that were kept under constant darkness after being entrained under 12-hour/12-hour light/dark cycle for 14 days (32). This specific dataset was selected due to the following reasons. First, *Bmal1* is the only gene that when knocked out in mice leads to a complete abolishment of circadian locomotor and feeding cycles, and as a result BMAL1 knockout mice is a widely-accepted model for circadian clock ablation (34). Secondly, in this study, BMAL1 knockout mice were kept in constant darkness, thus ruling out the possibility that any ultradian rhythms hereby uncovered are driven by light cues. Thirdly, of all the available temporal transcriptome dataset from BMAL1 knockout mice, it has the best temporal resolution (at 4-hour interval), duration (for a total of 48 hours) and sample size (n of four), thus permitting rigorous testing for ∼12-hour ultradian rhythms.

To unbiasedly reveal all oscillations from BMAL1 knockout mice liver, we initially applied the eigenvalue/pencil method (11, 12), which is a nonparametric spectrum analytical algorithm that deconvolutes raw temporal data into a linear combination of exponentials (sinusoidal waveforms with a decay factor) plus noise. Consistent with the role of BMAL1 as a master regulator of circadian rhythms, the prevailing population of circadian oscillations observed in wild-type mice are lost in the absence of BMAL1 (**Figure 1A**). By contrast, 3,589 genes cycling with periods between 10-13 hours were identified (**Figure 1A, B and Table S1**). Since the eigenvalue/pencil algorithm is a non-statistical signal processing method, we used a permutation-based method that randomly shuffles the time label of gene expression data to determine the false discovery rate (FDR) for the ∼12-hour genes (16, 33). This way, the FDR for the ∼12-hour genes was estimated to be 0.31, a range in line with prior studies of circadian rhythms (33, 35, 36). Furthermore, the calculated FDR is likely an overestimation since amplitude and phase information for the ∼12-hour rhythms is not considered in the permutation dataset.

**Figure 1.**
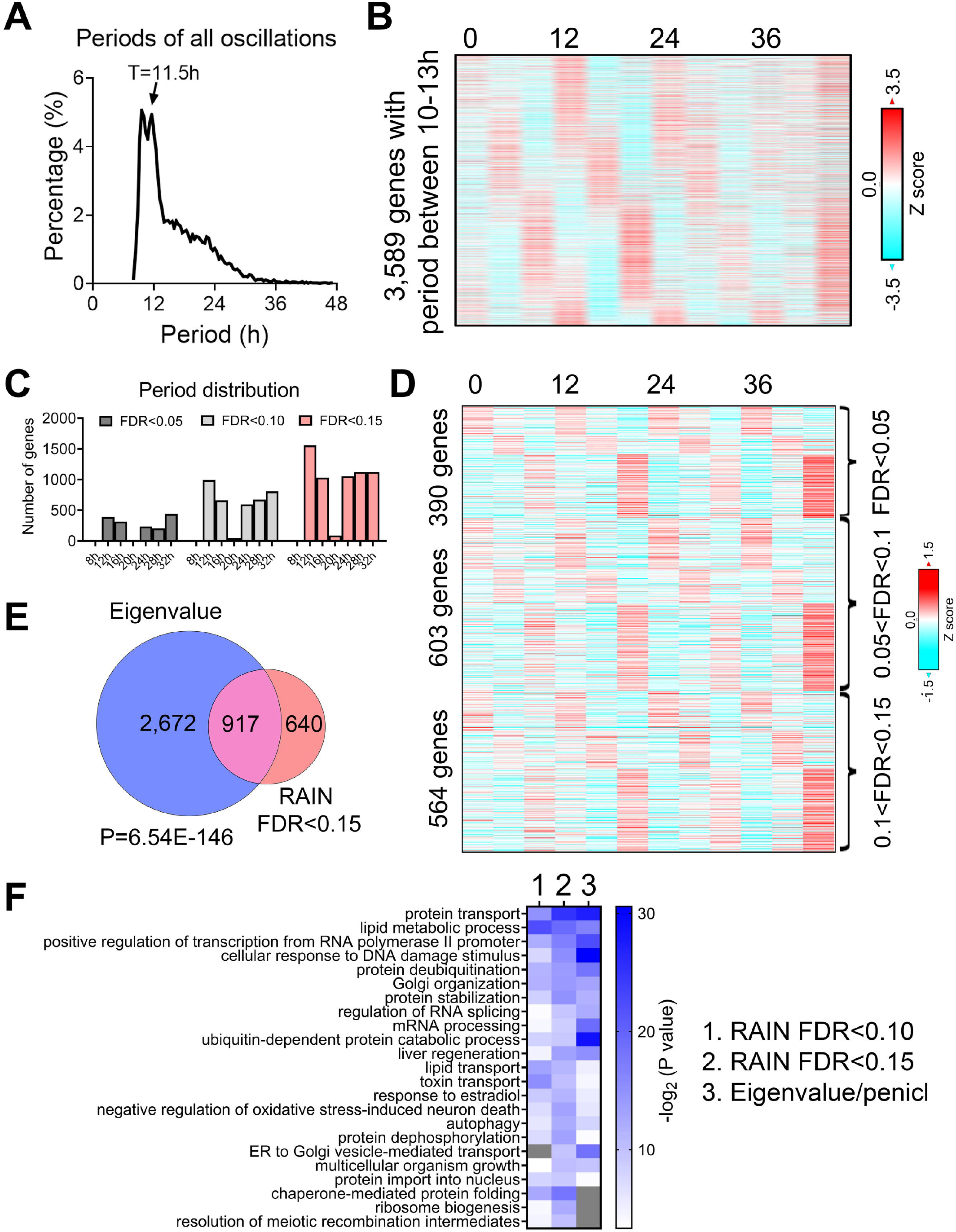
Prevalent ∼12-hour rhythms of gene expression in the liver of BMAL1 knockout mice. (**A**) Distribution of periods uncovered from all oscillations by the eigenvalue/pencil method. (**B**) Heatmap of ∼12-hour rhythms of gene expression uncovered by the eigenvalue/pencil method. (**C**) Number of genes cycling with different periods uncovered by the RAIN method, with different FDR cut-off. (**D**) Heatmap of ∼12-hour rhythms of gene expression ranked by different FDR cut-off. (**E**) Venn diagram comparing ∼12-hour transcriptome uncovered by the eigenvalue/pencil and RAIN (FDR cut-off of 0.15) method. P value calculated by Chi-square test. (**F**) GO analysis showing enriched biological pathways of ∼12-hour genes revealed by different methods using all mice genes as background.

To assess the robustness of the uncovered ∼12-hour oscillations in BMAL1 knockout mice, we applied an orthogonal rhythm-identification algorithm, RAIN (37), which detects rhythms with arbitrary waveforms exhibiting a single pre-specified period and has been successfully used to uncover ∼12-hour ultradian rhythms from high temporal resolution dataset in prior studies (15, 16). In line with the eigenvalue/pencil method, 12-hour were also the dominant oscillations identified by the RAIN method, with 390, 993 and 1,557 12-hour genes identified with FDR (or q value) cut-off of 0.05, 0.1 and 0.15, respectively (**Figures 1C, D, S1 A-C and Table S2**). Importantly, the robustness of 12-hour oscillations for genes under different FDR cut-off does not look visually different (**Figures 1D and S1 A, B**), and ∼12-hour genes identified by both methods further showed large convergence (**Figure 1E**).

We subsequently performed GO analysis on ∼12-hour genes identified by both methods using all mouse genes or all hepatically expressed genes as background, and identified commonly enriched pathways of lipid metabolism, proteostasis (including ribosome assembly, protein transport, protein processing, protein folding/stability/degradation, autophagy, and Golgi organization), and to a lesser degree, mRNA metabolism (transcription, RNA splicing and processing) (**Figures 1F and S1D**). Among these pathways, protein and mRNA metabolism were previously observed to be highly enriched in hepatic ∼12-hour transcriptome in wild-type mice as well (15, 16). Collectively, these results indicate the presence of prevalent ∼12-hour ultradian rhythms of gene expression in the absence of the circadian clock. Furthermore, since the locomotor activity and feeding behaviors of BMAL1 knockout mice are completely arrhythmic under constant darkness condition (34), it suggests that these ∼12-hour rhythms of gene expression are highly robust and resistant to feeding cues.

### Hepatic ∼12-hour rhythms of gene expression and functionality are conserved between wild-type and BMAL1 knockout mice under free-running condition

We next aim to determine the extent to which ∼12-hour transcriptome is conserved between wild-type and BMAL1 knockout mice. Those commonly found in both genotypes are likely controlled by the 12-hour oscillator, while “de novo” ∼12-hour genes only observed in BMAL1 knockout mice may reflect compensatory gain of new ultradian rhythms secondary to circadian clock ablation.

We selected the hepatic RNA-seq dataset published by Pan *et al* (16) for comparison due to the following reasons. First of all, this dataset has the highest temporal resolution (at 2-hour interval), duration (48 hours) and sample size (duplicates at each time point) among all temporal hepatic RNA-seq dataset for wild-type mice kept under constant darkness condition (38). Secondly, the strain (C57BL/6), gender (male) and age (8∼12 weeks) of these mice are comparable to those of BMAL1 knockout mice. Thirdly, comparison of data from mice housed in different locations further allow us to test for the robustness of ∼12-hour ultradian rhythms to vivarium housing environment variations.

Comparing ∼12-hour genes uncovered by the eigenvalue/pencil method in both datasets revealed a statistically significant (P value of 1.54e-6) overlap of 1,162 commonly shared genes, indicating preservation of ∼12-hour gene programs between wild-type and BMAL1 knockout mice (**Figures 2A, B and S2A**). The relative amplitude (absolute amplitude normalized to average gene expression) of shared ∼12-hour genes is comparable between the two genotypes, albeit with a slight (but statistically significant) increase observed in BMAL1 knockout mice (**Figure 2C**). By contrast, we did observe a difference in their phase distributions. Genes peaking around CT2 and CT14 in wild-type mice are phase-shifted to ∼CT8 and ∼CT20 (red rectangle in **Figure 2D**), and genes peaking around CT8 and CT20 in wild-type mice are shifted to peak around CT0 and CT12 (green circle in **Figure 2D**) in BMAL1 knockout mice.

**Figure 2.**
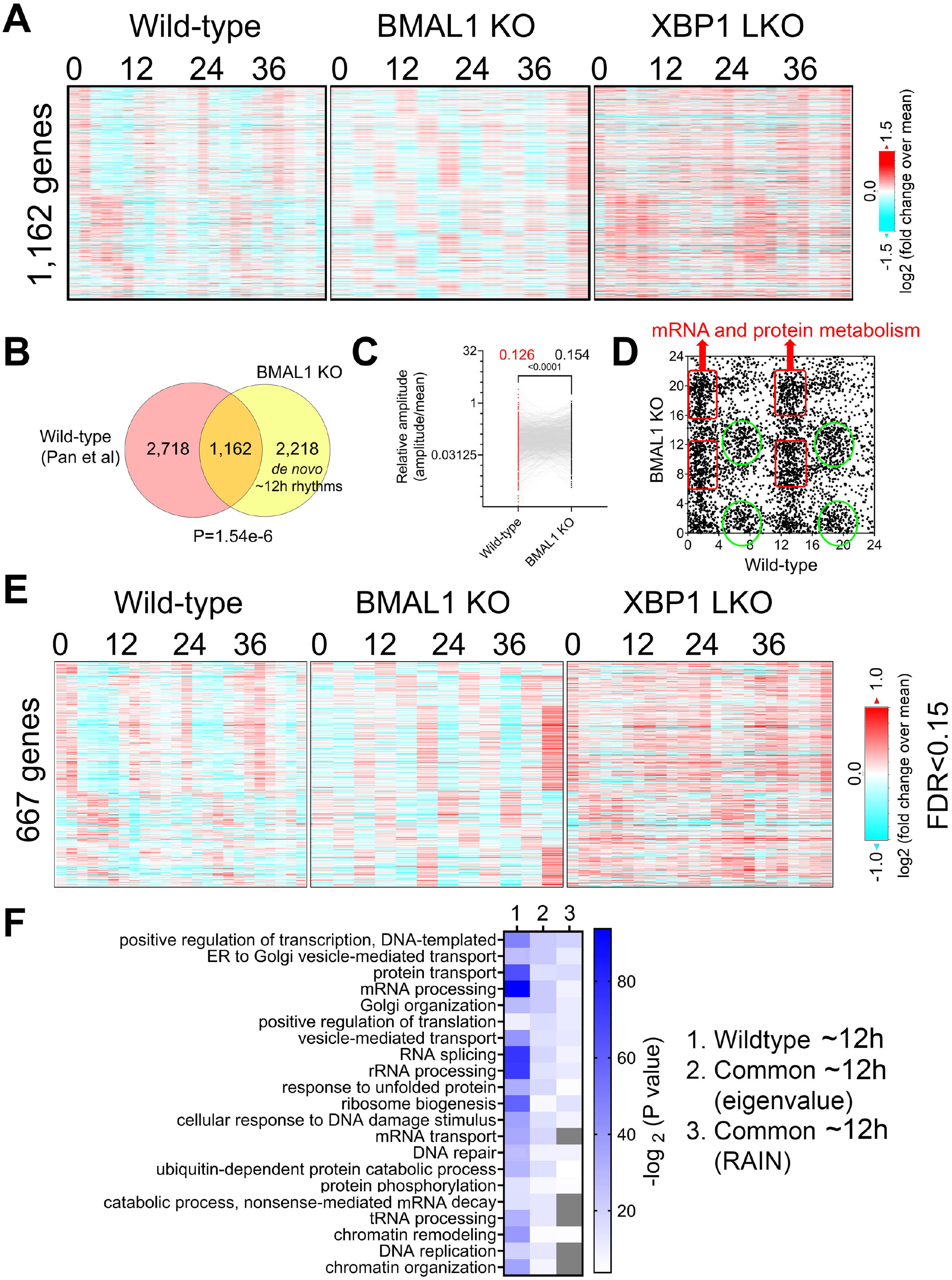
Preservation of ∼12-hour gene program and functionality between wild-type and BMAL1 knockout mice. (**A**) Heatmap of 1,162 commonly found ∼12-hour rhythm gene expression in wild-type, BMAL1 knockout and XBP1 LKO mice. (**B**) Venn diagram comparing ∼12-hour rhythm genes in wild-type and BMAL1 knockout mice. P value is calculated by Chi-square test. (**C**) Relative amplitude of ∼12-hour oscillations for 1,162 genes in wild-type and BMAL1 knockout mice. (**D**) Scatter plot comparing the phases of ∼12-hour oscillations in wild-type and BMAL1 knockout mice. **Genes within red rectangle are enriched in mRNA and protein metabolism**. (**E**) Heatmap of 667 commonly found ∼12-hour rhythm gene expression in wild-type, BMAL1 knockout and XBP1 LKO mice (FDR cut-off of 0.15). (**F**) GO analysis showing enriched biological pathways of ∼12-hour genes found in wild-type or in both wild-type and BMAL1 knockout mice using all mice genes as background.

GO terms (regardless of the background gene sets used) associated with those 1,162 commonly shared ∼12-hour genes are highly similar to those of ∼12-hour genes in wild-type mice, namely, proteostasis (protein transport, Golgi organization, translation regulation, UPR, tRNA processing, and ubiquitin-mediated protein degradation) and mRNA metabolism (transcription regulation, mRNA processing and splicing and mRNA transport) (**Figures 2F and S2 A, B**). Further examining ∼12-hour rhythms stratified by phases revealed that genes peaking between CT0 and CT4 (and between CT12 and CT16) in wild-type mice and shifting to between CT6 and CT12 (and between CT18 and CT24) in BMAL1 knockout mice are very strongly enriched in the same pathways of mRNA and protein metabolism, while genes peaking between CT4 to CT10 (and between CT16 and CT22) in wild-type mice and shifting to between CT10 and CT15 (and between CT22 and CT27) in BMAL1 knockout mice are very modestly enriched in pathways of GTPase activity, cell polarity and amino acid metabolism (**Figures 2D and S2C**). The convergence of ∼12-hour genes and their functionality between wild-type and BMAL1 knockout mice are further validated with the RAIN method (FDR cut-off of 0.15) (**Figures 2E, F and S2A, B**).

In addition to commonly shared ∼12-hour genes, many genes with ∼12-hour rhythms exclusively identified in either wild-type or BMAL1 knockout mice were present (**Figure 2B**). A close examination of their period change in the opposite genotype reveals that 2,718 ∼12-hour genes only identified in wild-type mice often exhibit periods slightly lower than 10-hour in BMAL1 knockout mice, thus being labelled as not having ∼12-hour oscillations (**Figure S3A**). These genes are also enriched in protein and mRNA metabolic pathway as expected regardless of the background gene sets used (**Figure S3B, C**). On the contrary, many of the 2,218 ∼12-hour genes exclusively observed in BMAL1 knockout mice exhibit circadian period in wild-type mice and these genes are strongly enriched in lipid metabolism, mitosis, complement activation and Notch signaling, pathways that are not strongly associated with ∼12-hour transcriptome in wild-type mice (**Figure S3A-C**). The latter group can be categorized as *de novo* ∼12-hour genes that are gained in BMAL1 knockout mice secondary to the ablation of circadian clock. Taken together, these findings provide additional evidence to support the existence of a 12-hour oscillator that controls ∼12-hour rhythms of gene expression of protein and mRNA metabolism in mice liver.

### Many hepatic genes are under dual circadian clock and 12-hour oscillator control

∼12-hour genes commonly identified in both wild-type and BMAL1 knockout mice are likely under 12-hour oscillator control. In agreement with this hypothesis, their ∼12-hour rhythms of gene expression are impaired in XBP1 LKO mice (16) (**Figure 2A and 2E**). Conversely, as previously reported, core circadian clock and the majority of circadian output genes are maintained in XBP1 LKO mice (15, 16), and abolished in BMAL1 knockout mice, as expected (**Figure 3A**). While the repertoire and functionality of circadian and ∼12-hour genes in wild-type mice are largely separate (**Figures 3B, C and S4A**), we did observe hundreds of genes having both circadian and ∼12-hour rhythms superimposed (**Figure 3C**). The phenomenon of superimposition of circadian and ultradian rhythms in the same genes are widespread and have been previously reported in many different organisms by other groups (13, 39, 40). Using a very stringent criterion where the superimposed circadian and ∼12-hour rhythms need to be abolished in BMAL1 KO and XBP1 LKO mice, respectively, but maintained in XBP1 LKO and BMAL1 KO mice, respectively, we identified 141 genes under robust dual control of BMAL1-dependent circadian clock and XBP1s-dependent 12-hour oscillator (**Figure 3C and Table S3**). Examples of such genes include chaperone *Cct3*, mitochondria import receptor *Tomm40l* (Tom40b), master transcriptional regulator of fatty acid oxidation *Ppara*, and RNA helicase *Ddx1* (**Figures 3D and S4B**). In all cases, eigenvalue/pencil method revealed coexistence/superimposition of both circadian and ∼12-hour rhythms (sometimes with additional ∼4.8-hour or ∼8-hour rhythms as well) in wild-type mice, and more importantly, the linear addition of these rhythms closely matches the raw temporal gene expression profiles, indicating the robustness of deconvolution (**Figures 3D and S4B**). Interestingly, while in BMAL1 KO and XBP1 LKO mice, circadian and ∼12-hour rhythms are no longer identified by the eigenvalue/pencil method, respectively, ultradian rhythms with even smaller periods (4.8∼8-hour) are often observed (**Figures 3D and S4B**), indicating the likely existence of additional oscillators for high frequency rhythms.

**Figure 3.**
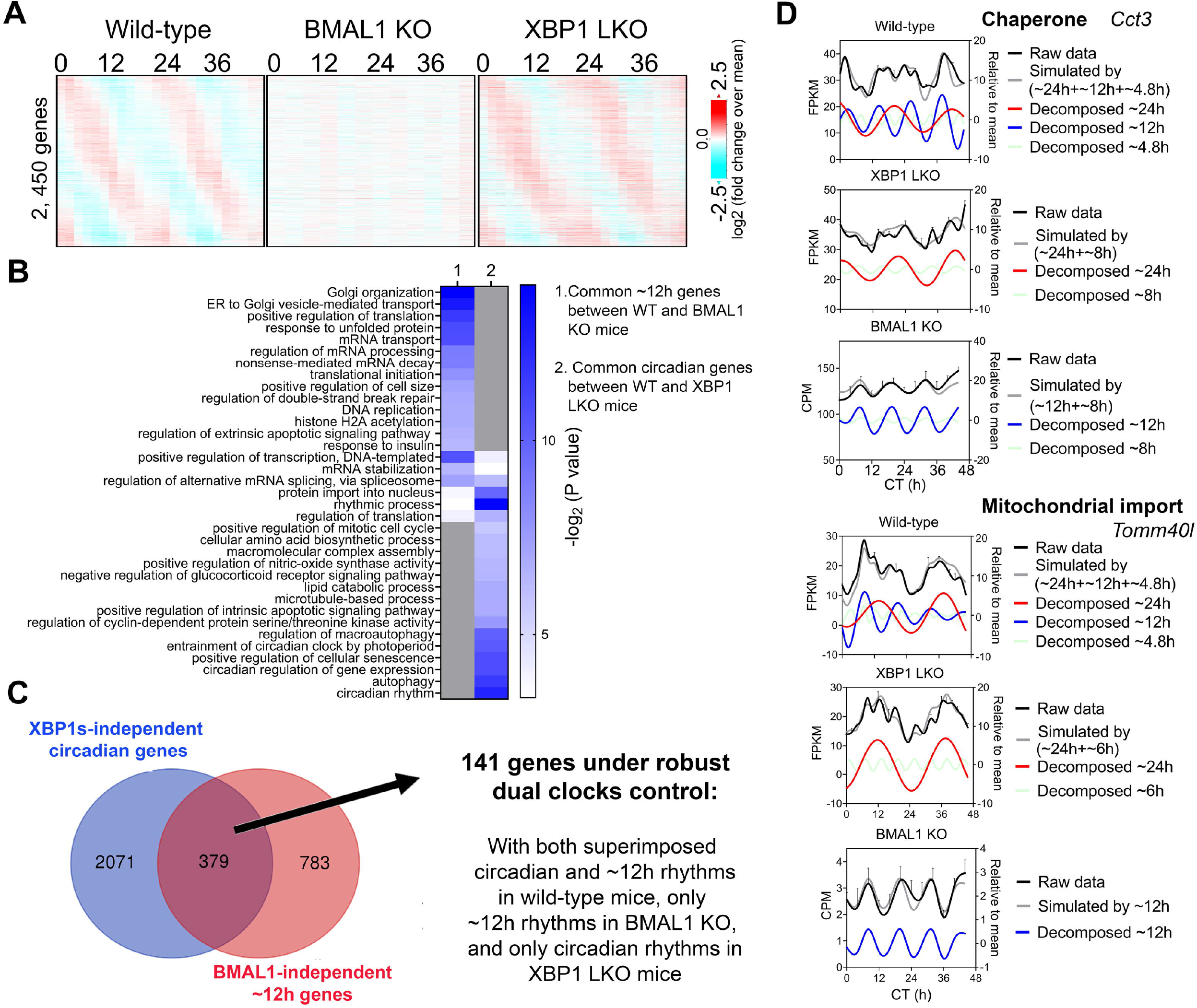
Many hepatic genes are under dual circadian clock and 12-hour oscillator control. (**A**) Heatmap of 2,450 commonly found circadian gene expression (common between wild-type and XBP1 LKO mice) in wild-type, BMAL1 knockout and XBP1 LKO mice. (**B**) GO analysis showing enriched biological pathways of common ∼12-hour (between wild-type and BMAL1 knockout mice) and circadian (between wild-type and XBP1 LKO mice) using hepatically expressed genes as background. (**C**) Venn diagram illustrating a total of 141 genes under robust dual clock control. (**D**) Eigenvalue/pencil deconvolution of *Cct3* and *Tomm40l* temporal gene expression profiles in wild-type, BMAL1 knockout and XBP1 LKO mice. Gray line in each graph illustrates simulated temporal gene expression profile by addition of all superimposed oscillations.

### ELF1, KLF7 and ATF6B are novel putative transcription regulators of the hepatic 12-hour oscillator

To infer the gene regulatory network governing ∼12-hour oscillator, we performed motif analysis on the commonly shared ∼12-hour genes between wild-type and BMAL1 knockout mice identified by either the eigenvalue/pencil or RAIN methods. Regardless of the method used, we identified top motifs associated with KLF, ETS, NFY and basic leucine zipper (including ATF6, XBP1, CREB3, CREB3L2 and CREBRF) families of transcription factors (**Figure 4A**), consistent with previous results (16, 17). *Xbp1* itself exhibited a ∼13-hour oscillation in the liver of BMAL1 knockout mice (**Figure 4B**). We further cross-referenced enriched motifs with transcription factors exhibiting ∼12-hour rhythms in both wild-type and BMAL1 knockout mice, but not in XBP1 LKO mice, and identified KLF7, ELF1 and ATF6B as strong putative transcriptional regulators of 12-hour oscillator. Interestingly, *KLF7* was previously shown to also exhibit a ∼12-hour rhythm of expression in human dorsolateral prefrontal cortex (27). In addition, we further identified NYFB and CREBRF as two transcriptional factors exhibiting ∼12-hour rhythms in all three genotypes of mice (**Figure 4C**). Of these transcription factors, hepatic XBP1s, NFYA, NFYC and GABPB1 expressions further exhibit ∼12-hour rhythms at the protein level in wild-type mice, as previously measured (41) and quantified (20), while the expression levels of GABPA/B2, ATF6/6B, NFYB, KLF7 and ELF1 were not reported from the same study.

**Figure 4.**
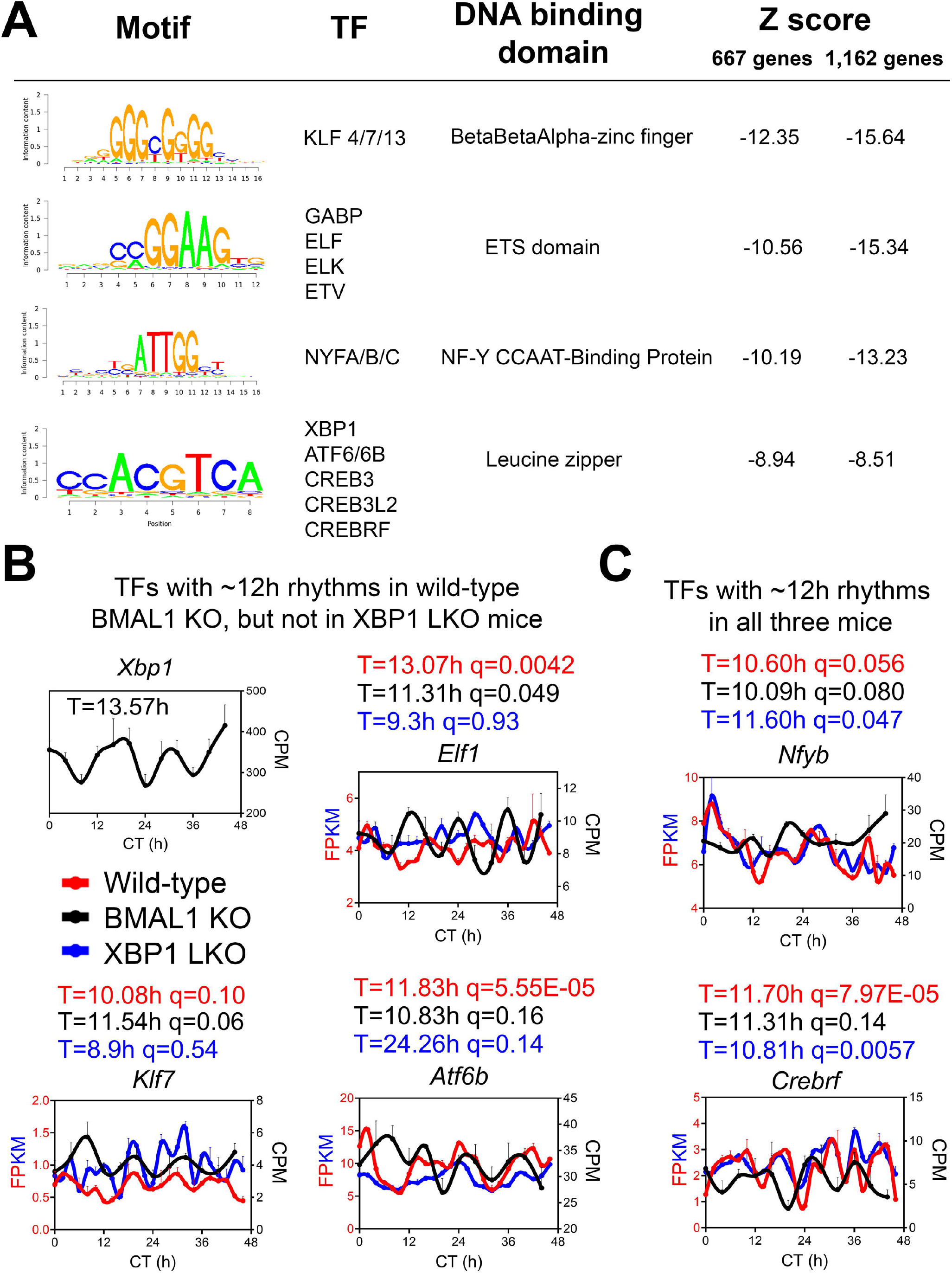
ELF1, KLF7 and ATF6B are putative transcription regulators of 12-hour oscillator. (**A**) Motif analysis of promoter regions of common ∼12-hour rhythm genes identified in wild-type and BMAL1 knockout mice by either method. (**B, C**) Expression of different transcription factors in wild-type, BMAL1 knockout and XBP1 LKO mice. Period identified by the eigenvalue/pencil method, and q value (FDR) for having 12-hour rhythms of gene expression via the RAIN method were shown for each gene in each genotype.

To ensure that our analysis is robust to datasets used, we further compared the ∼12-hour hepatic transcriptome of BMAL1 knockout mice with an additional wild-type dataset published by the same group (32). The relatively low resolution (4-hour interval for 48-hours under constant darkness) renders this dataset not ideal for ultradian rhythms detection by statistical methods like RAIN due to the superimposition of strong circadian rhythms for many genes. Nonetheless, using eigenvalue/pencil method, we uncovered hundreds of overlapping ∼12-hour transcriptome between wild-type and BMAL1 knockout mice that has 1) similar amplitude but distinct phases, 2) strong enrichment in proteostasis pathways, and 3) over-representation of KLF, bZIP, ETS or NYF TFs motifs in their promoters (**Figure S5A-H**), all consistent with the results described above. Collectively, our analysis of ∼12-hour rhythms in wild-type, BMAL1 knockout and XBP1 LKO mice provided novel mechanistic insights on the regulation of mammalian 12-hour oscillator.

### Cell-autonomous ∼12-hour rhythms of gene expression involved in mRNA and protein homeostasis are present in *Drosophila* S2 cells that lack canonical circadian clock gene expression

Having established the preservation of ∼12-hour rhythms of gene expression in the absence of circadian clock in mice, we wanted to further corroborate this finding in additional animals that also lack a canonical circadian clock. We focused upon the fruit fly *Drosophila melanogaster*, where ∼12-hour rhythms, to the best of our knowledge, have never been reported before. In *Drosophila melanogaster*, the model of the circadian clock regulation involves transcription factors CYCLE (CYC) and CLOCK (CLK), the homologs of BMAL1 and CLOCK in mammals, which in turn control the transcription of several repressive clock genes including *period* (*per*) and *timeless* (*tim*) (42). Besides a subset of cells located in the principal pacemaker driving daily activity rhythms and physiology, most other cells including *Drosophila* Schneider 2 (S2) cells [originally derived from a primary culture of late-stage embryos (43)] do not express these core circadian clock genes and therefore do not have the necessary apparatus to form a TTFL to drive circadian oscillations (33, 44, 45). We thus deduce that if ∼12-hour rhythms of gene expression with functionality like those identified in mice can be observed in S2 cells, it will provide additional evidence for the existence of a cell-autonomous 12-hour oscillator independent from the canonical circadian clock.

We analyzed a recently published temporal RNA-seq data (33) collected from temperature entrained S2 cells (2 days of temperature cycles with 12-hour at 28°C, 12-hour at 23°C before free-running at 28°C) at 3-hour interval for a total duration of 60 hours. Using both eigenvalue/pencil and RAIN methods, we identified tens of hundreds of transcripts cycling with an ultradian period between 11 and 15 hours (**Figure 5A-D, Table S4 and S5**). This period range is longer than what was observed in mice, likely due to the likewise longer circadian period (22∼28 hours) identified in S2 cells (**Figure 5A-C**). This way, both ∼12 and ∼24-hour rhythms are still cycling at harmonics frequencies with each other in S2 cells. GO analysis of these ∼12-hour ultradian genes revealed top enriched pathways of protein transport (such as *Rab1*) and mRNA splicing (such as *Pnn*), distinct from those associated with circadian rhythms but similar to those associated with ∼12-hour genes in mice (**Figures 5E-F, and S6**). Among the putative/confirmed transcription regulators governing ∼12-hour rhythms in mice, we found the fly orthologs of *Gabpa, Son, Atf6b* and *Elf1* exhibiting a period between 11 and 14 hours in S2 cells (**Figure 6**). The fly orthologs of *Nfya* and *Xbp1* exhibit a longer period of 15∼16 hour in S2 cells (**Figure 6**). In sum, we herein provided additional evidence supporting the existence of ∼12-hour rhythms of gene expression of mRNA and protein metabolism in clock-less fly cells. More importantly, the fact that ∼12-hour rhythms of gene expression implicated in protein and mRNA metabolism have been found in divergent species of sea anemone (46), *C. elegans* (47), fly, mice and human (27) argue for the strong evolutionary conservation of the ∼12-hour oscillator.

**Figure 5.**
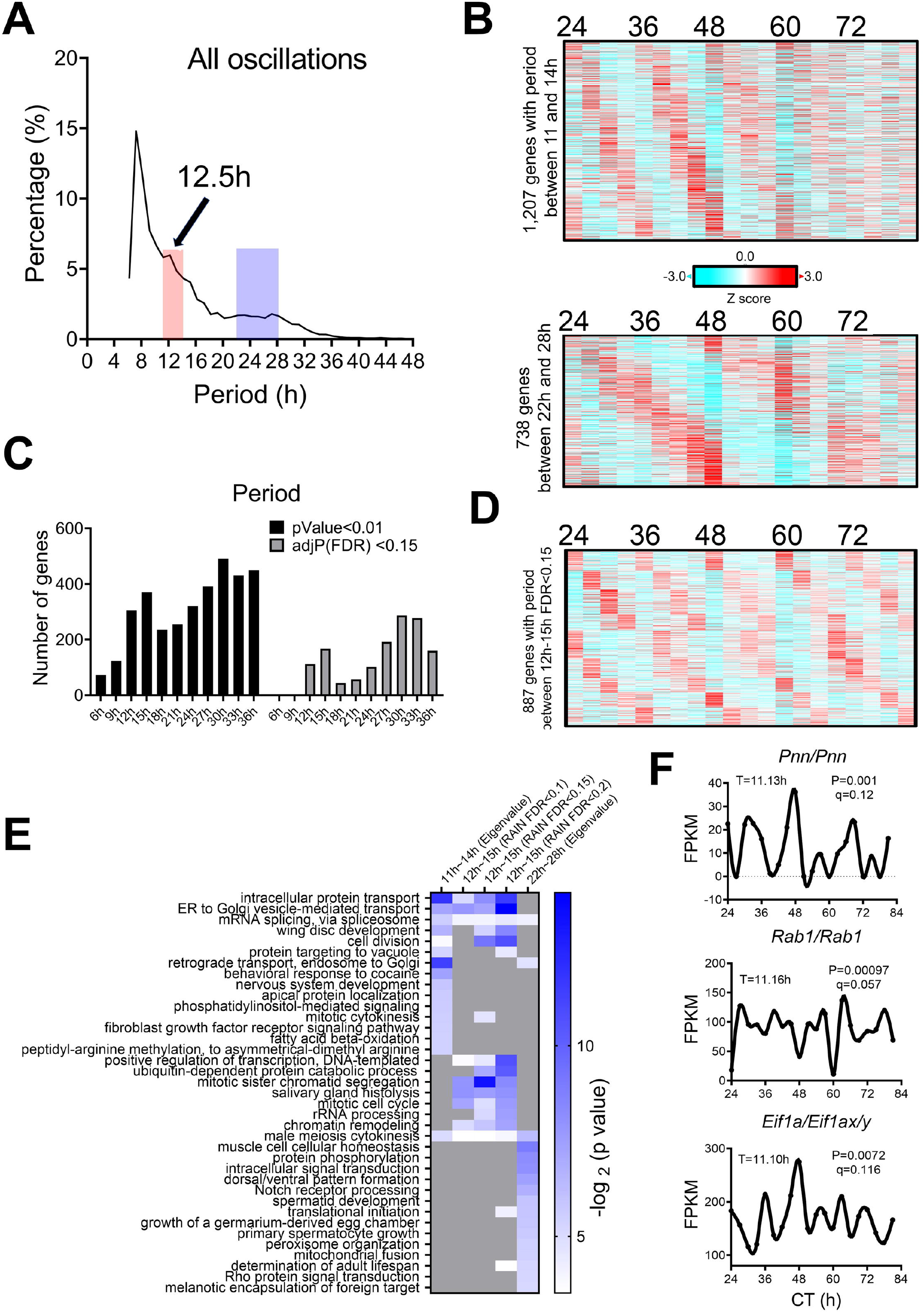
∼12-hour rhythms of gene expression are present in S2 cells. (**A**) Distribution of periods uncovered from all oscillations by the eigenvalue/pencil method. (**B**) Heatmap of ∼12-hour (top) and circadian rhythms (bottom) of gene expression uncovered by the eigenvalue/pencil method. (**C**) Number of genes cycling with different periods uncovered by the RAIN method, by p value or FDR cut-off. (**D**) Heatmap of ∼12-hour rhythms of gene expression with FDR cut-off of 0.15 via RAIN. (**E**) GO analysis of ∼12-hour and circadian rhythms uncovered by different methods/FDR cut-off in S2 cells using all fly genes as background. (**F**) Representative expression of selective genes. Period identified by the eigenvalue/pencil method, and q value (FDR) for having ∼12-hour rhythms of gene expression via the RAIN method were shown for each gene.

**Figure 6.**
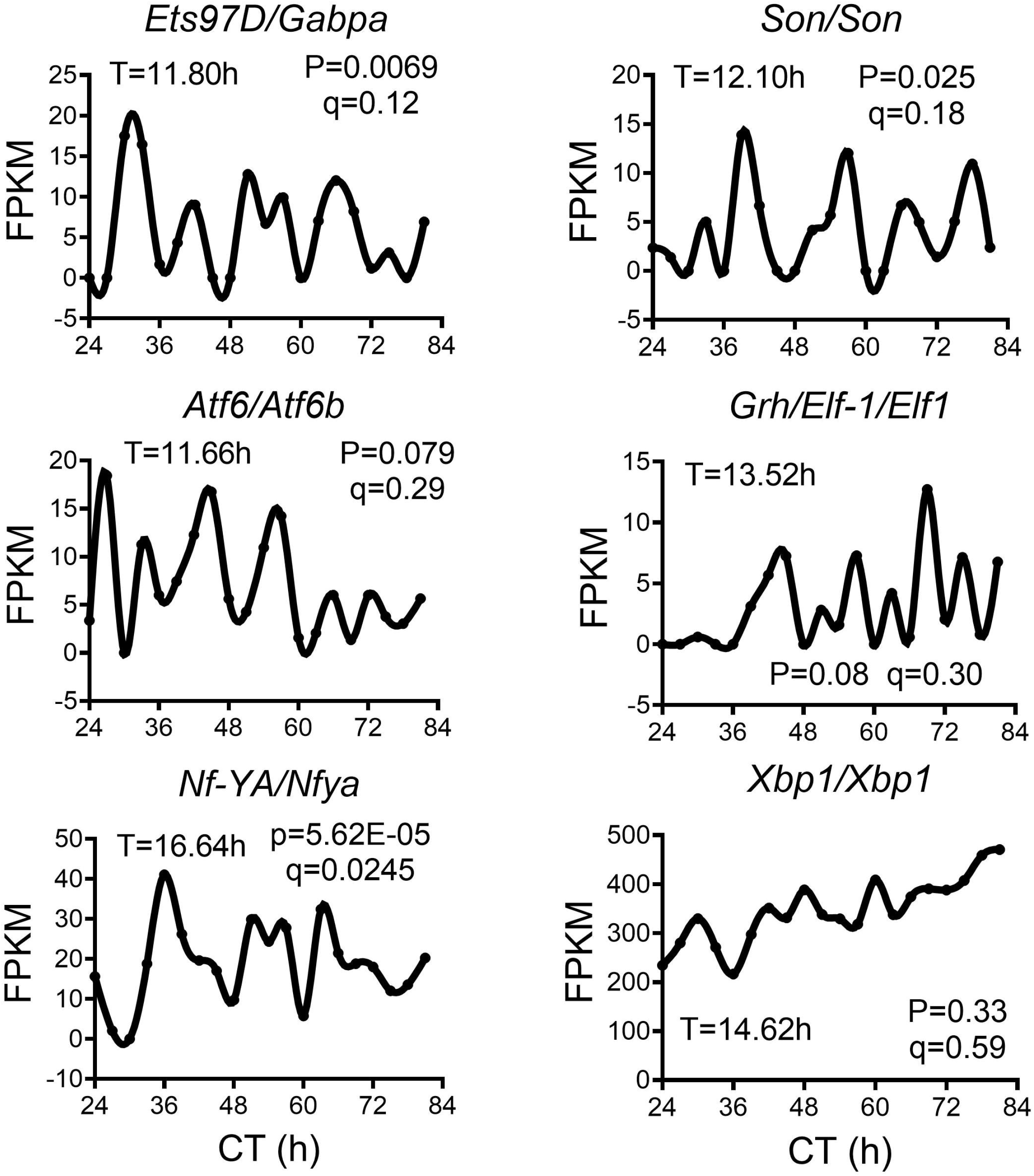
∼12-hour rhythms of transcription regulators in S2 cell. Representative expression of selective transcription regulators in S2 cell. Period identified by the eigenvalue/pencil method, and q value (FDR) for having ∼12-hour rhythms of gene expression via the RAIN method were shown for each gene.

## Discussion

In this study, by using two different analytical methods eigenvalue/pencil and RAIN, we uncovered prevalent ∼12-hour rhythms of gene expression in two different models of circadian clock deficiency: BMAL1 knockout mice and *Drosophila* S2 cells. In both models, the ∼12-hour gene programs are strongly enriched in protein and mRNA metabolism, consist with the gene signature previously identified for ∼12-hour/circatidal genes from wild-type mice, sea anemone and even *C. elegans* (18). These findings thus lend further support for the 12-hour oscillator hypothesis on the origin and regulation of ∼12-hour ultradian rhythms.

In an early study (28), Cretenet et al., reported that in the liver of CRY1/CRY2 double knockout mice, ∼12-hour rhythms of expression of several UPR genes (including *Xbp1* itself) were disrupted. We believe that at least three factors can account for the discrepancies observed between BMAL1 knockout and CRY1/CRY2 knockout mice. First of all, due to the limit of technologies available at the time, the complete hepatic temporal transcriptome of CRY1/CRY2 knockout mice were not profiled. Thus, there remains the possibility that many ∼12-hour rhythms of gene expression are also maintained in the liver of CRY1/CRY2 knockout mice, and future high temporal resolution profiling from these mice is needed to answer this question. Secondly, since the 12-hour rhythms are sensitive to heightened metabolic and ER stress (11, 16, 28), it is highly likely that the observed changes of 12-hour rhythms in Cry1/Cry2 knockout mice is a consequence of the metabolic deficits observed in these mice, rather than due to the absence of a functional circadian clock per se. In fact, as reported in this study and stated by the authors (28), CRY1/CRY2 knockout mice liver exhibited a much higher level of triglycerides, which in turn can lead to chronic ER stress, and subsequently constitutive action of IRE1α/XBP1s signaling and impairment of ∼12-hour ultradian rhythms of gene expression of proteostasis. By contrast, BMAL1 knockout mice has a normal level of hepatic triglyceride and even increased insulin sensitivity compared to wild-type mice (48). As a result, hepatic ∼12-hour rhythms of gene expression are much better maintained in BMAL1 knockout mice than CRY1/CRY2 knockout mice. Finally, CRY proteins may impact overall proteostasis independently from its canonical function in regulating circadian rhythm, via protecting against proteome imbalance and proteotoxic stress as recently demonstrated (49).

While the period and amplitude of hepatic ∼12-hour rhythms of gene expression are not affected by BMAL1 ablation, we did observe a major shift in their phase distributions. In wild-type mice, genes involved in protein and mRNA metabolism peak around CT2 and CT14, corresponding to dawn and dusk (15, 16). By contrast, in the liver of BMAL1 knockout mice, many of their phases are shifted to CT8 and CT20. Overall, a decline of phase coherence of ∼12-hour rhythms was observed in BMAL1 knockout mice. Since the phases of 12-hour rhythms of gene expression in MEFs are not altered in the presence of transient BMAL1 knocking down *in vitro (11, 16)*, this phase decoherence may be attributed to the differential decoupling between different ∼12-hour genes and environmental input as a result of altered behaviors of BMAL1 knockout mice.

The conservation of ∼12-hour gene programs and functionality among Cnidarian, nematode, insects, and mammals argues for an ancient evolutionary origin of ∼12-hour oscillator. Since all life originated from the ocean and all in-land animals had to experience tidal environment during ocean-to-land transition in the early era of evolution, it is reasonable to hypothesize that mammalian 12-hour oscillator evolved from the circatidal clock of coastal and estuarine animals (7, 11, 16). Although the mechanisms underlying the marine circatidal rhythms remain an open field of research, more and more studies favor the circatidal clock hypothesis (50). For instance, in the crustacean *E. pulchra*, the 12-hour circatidal clock is dissociated from the circadian timekeeping system (51). In circatidal mangrove crickets, 12-hour rhythms persist in constant darkness, even after the removal of the optic lobe. Furthermore, these 12-hour rhythms are intact even when expression of components of the circadian TTFL such as *Clock* or *per* are reduced through siRNA-mediated knockdown (52-54). What adaptive advantage might 12-hour rhythms of gene expression confer? We reason that the ancient ∼12-hour rhythms that evolved in marine species in response to tidal cues have been coopted by mammals as an adaptive response to accommodate physiological transitions related to feeding, physical activity, and sleep that are temporally concentrated at dawn and dusk.

Last but not the least, while our study provides strong support for the ∼12-hour oscillator hypothesis, we are not ruling out the other two possibilities, namely, that some ∼12-hour rhythms may not be cell-autonomous and can be controlled by a combination of the circadian clock and environmental cues, or that they can be regulated by two anti-phase circadian transcriptional factors in a cell-autonomous manner. Future studies using additional *in vivo* and *in vitro* models of circadian deficiency are needed to fully understand the complex relationship between ultradian and circadian rhythms (19).

## Materials and methods

### Identification of the oscillating transcriptome

For BMAL1 knockout or WT mice RNA-seq data (GSE171975), raw counts were first normalized to total sequencing depth to get counts per millions (CPM) values, which were used for cycling transcripts identification by either the eigenvalue/pencil or RAIN methods. For the eigenvalue/pencil method (11, 12), the mean expression at each time point were calculated and used as input. A maximum of two oscillations were identified for each gene. Criterion for ∼12-hour genes: period between 10 hour to 13-hour, decay rate between 0.8 and 1.2. The relative amplitude was calculated by dividing the amplitude by the mean expression value for each gene. To determine FDR, we used a permutation-based method that randomly shuffles the time label of gene expression data and subjected each permutation dataset to the eigenvalue/pencil method applied with the same criterion (33). These permutation tests were run 5,000 times, and FDR was estimated by taking the ratio between the mean number of rhythmic profiles identified in the permutated samples (false positives) and the number of rhythmic profiles identified in the original data. Analyses were performed in MatlabR2017A. RAIN analysis was performed as previously described in Bioconductor (3.4) (http://www.bioconductor.org/packages/release/bioc/html/rain.html) (37). FDR (also known as q value) was calculated using the Benjamini-Hochberg procedure.

For RNA-seq in S2 cells (GSE102495), FPKM values reported in the original study were used for cycling transcripts identification. Eigenvalue/pencil analysis was performed as previously described except for that genes with periods between 11 and 14-hour are defined as ∼12-hour genes. For RAIN analysis, the FPKM data was first subject to polynomial detrend (n=2). For both datasets, heat maps were generated with Gene Cluster 3.0 and TreeView 3.0 alpha 3.0 using either Z score or log2 (fold change over mean) values.

### Gene ontology analysis

DAVID (Version 2021) (55) (https://david.ncifcrf.gov) was used to perform Gene Ontology analyses. Briefly, gene names were first converted to DAVID-recognizable IDs using Gene Accession Conversion Tool. The updated gene list was then subject to GO analysis using either all *Mus musculus* (for mice) or *Drosophila melanogaster* (for S2 cells) genes or hepatically expressed genes (for mice) or all genes expressed in S2 cells as background and with Functional Annotation Chart function. GO_BP_DIRECT was used as GO categories. Only GO terms with a p value less than 0.05 were included for further analysis.

### Motif analysis

Motif analysis was performed with the SeqPos motif tool (version 0.590) embedded in Galaxy Cistrome using all motifs within the mouse reference genome mm9 as background. Genome region spanning 0.5kb upstream and 05kb downstream of transcription start were used as input.

### Statistical Analysis

Data were analyzed and presented with GraphPad Prism software. Plots show individual data points and bars at the mean and ± the standard error of the mean (SEM). One-tailed t-tests were used to compare means between groups, with significance set at p < 0.05.

## Author contributions

Conceptualization, B.Z; Methodology, B.Z.; Investigation, B.Z., and S.L. Writing – Original Draft, B.Z; Writing – Review & Editing, all authors; Funding Acquisition, B.Z., and S.L. Resources, B.Z; Supervision, B.Z.

## Acknowledgements

B. Zhu was supported by grants 1DP2GM140924 and 1R21AG071893 through the NIH. This research was supported in part by the University of Pittsburgh Center for Research Computing through the resources provided. Specifically, this work used the HTC cluster, which is supported by NIH award number S10OD028483. This research project was supported in part by the Pittsburgh Liver Research Centre supported by NIH/NIDDK Digestive Disease Research Core Center grant P30DK120531.

