## Supplemental materials for "Preservation of ∼12-hour ultradian rhythms of gene expression of mRNA and protein metabolism in the absence of canonical circadian clock"

### Supplemental figures:

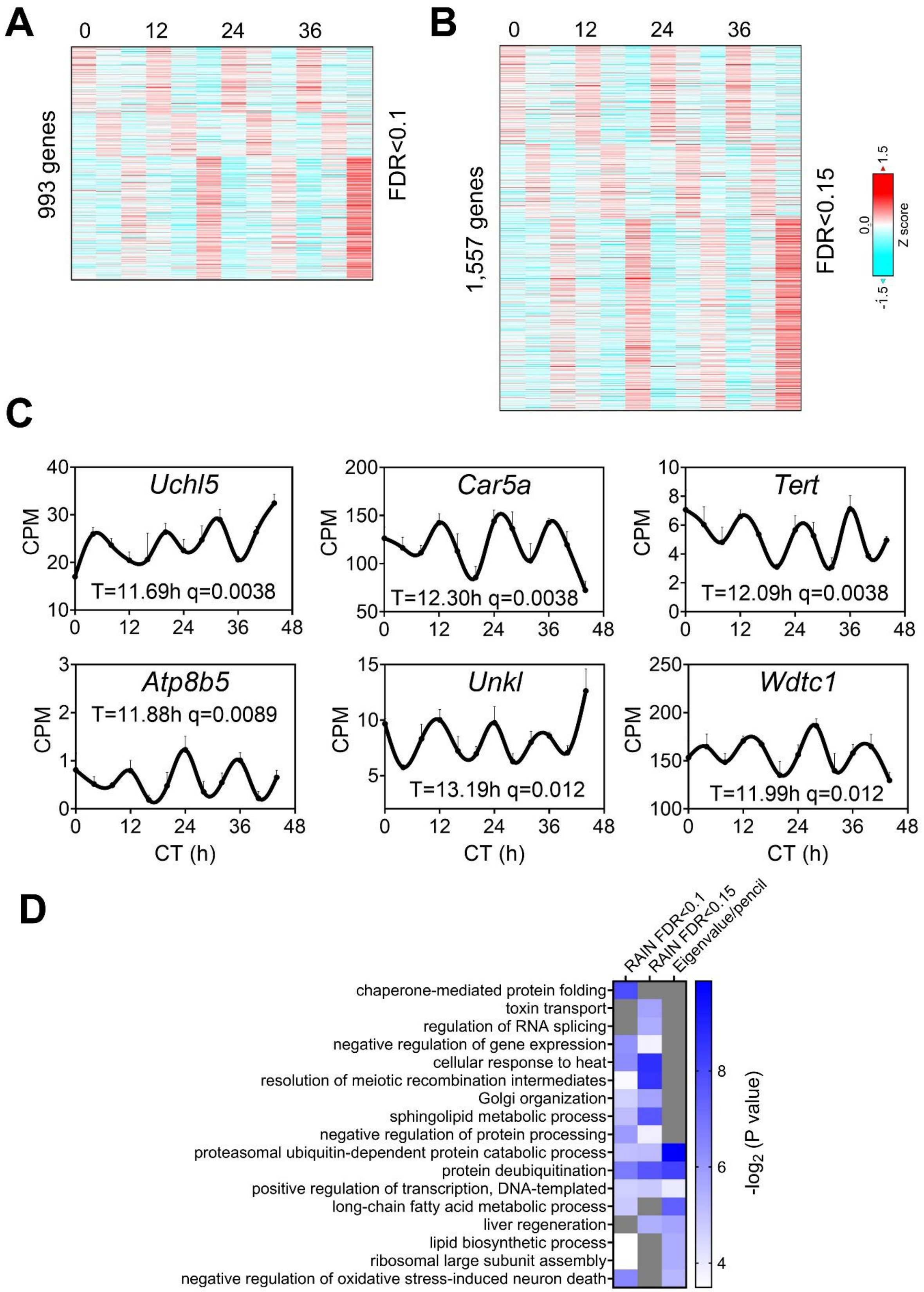
A
0
12
24
36
993 genes
FDR<0.1
B
0
12
24
36
1,557 genes
FDR<0.15
1.5
0.0
-1.5
Z score
C
Uchl5
T=11.69h q=0.0038
Car5a
T=12.30h q=0.0038
Tert
T=12.09h q=0.0038
Atp8b5
T=11.88h q=0.0089
Unkl
T=13.19h q=0.012
Wdtc1
T=11.99h q=0.012
CPM
CT (h)
D
RAIN FDR<0.1
RAIN FDR<0.15
Eigenvalue/pencil
chaperone-mediated protein folding
toxin transport
regulation of RNA splicing
negative regulation of gene expression
cellular response to heat
resolution of meiotic recombination intermediates
Golgi organization
sphingolipid metabolic process
negative regulation of protein processing
proteasomal ubiquitin-dependent protein catabolic process
protein deubiquitination
positive regulation of transcription, DNA-templated
long-chain fatty acid metabolic process
liver regeneration
lipid biosynthetic process
ribosomal large subunit assembly
negative regulation of oxidative stress-induced neuron death
-log2 (P value)

**Figure S1. Prevalent ~12-hour rhythms of gene expression in the liver of BMAL1 knockout mice.**

(**A, B**) Heatmap of 12-hour rhythms of gene expression uncovered by RAIN with FDR cut-off of 0.1 (**A**) or 0.15 (**B**). (**C**) Expression of six genes with the lowest q values uncovered by RAIN in BMAL1 knockout mice. Period identified by the eigenvalue/pencil method, and q value (FDR) for having 12-hour rhythms of gene expression via the RAIN method were shown for each gene. (**D**) GO analysis showing enriched biological pathways of ~12-hour genes revealed by different methods using hepatically expressed genes as background.

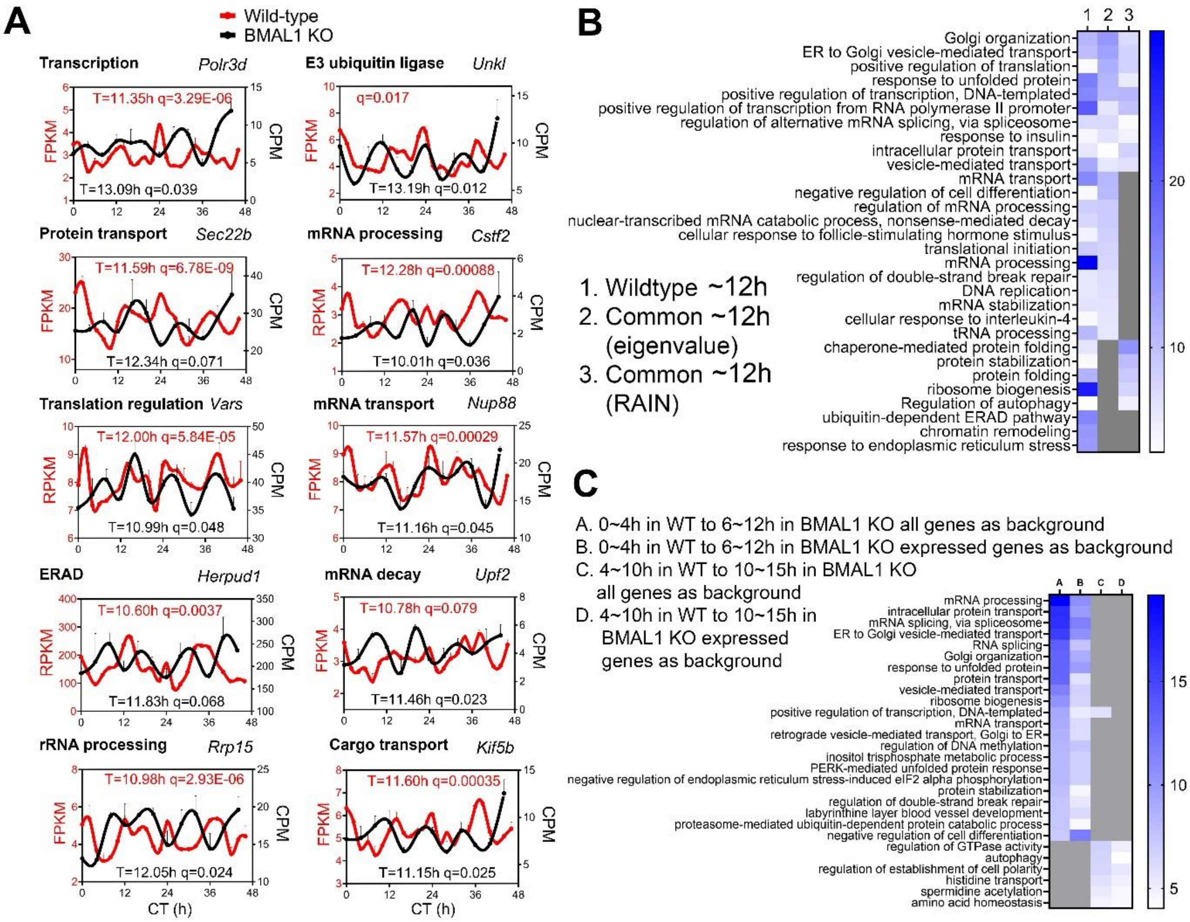


**Figure S2. Preservation of ~12-hour gene programs and functionality in wild-type and BMAL1 knockout mice.**

(**A**) Representative ~12-hour genes expression in both wild-type and BMAL1 knockout mice. Period identified by the eigenvalue/pencil method, and q value (FDR) for having 12-hour rhythms of gene expression via the RAIN method were shown for each gene in each genotype. (**B**) GO analysis showing enriched biological pathways of ~12-hour genes found in wild-type or in both wild-type and BMAL1 knockout mice using all hepatically expressed genes as background. (**C**) GO analysis showing enriched biological pathways of ~12-hour genes stratified by phases using all mice genes or hepatically expressed genes as background.

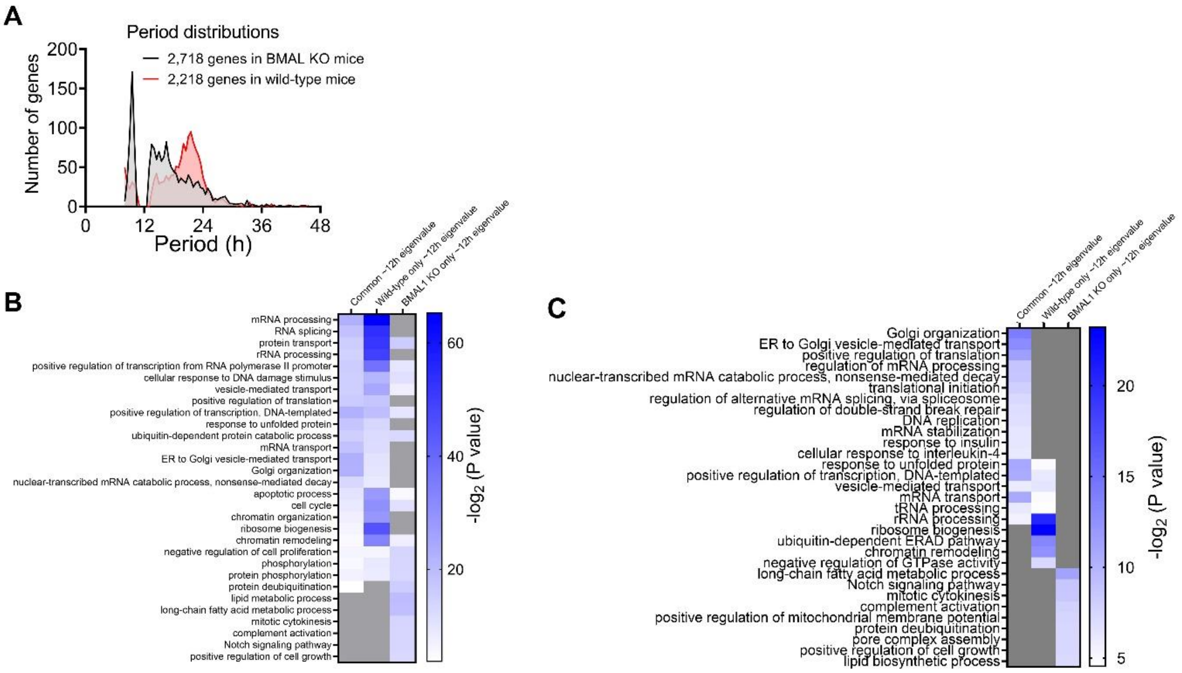


**Figure S3. Distinct functionality of shared and BMAL1 knockout-specific ~12-hour rhythms of gene expression.**

(**A**) Distribution of periods of wild-type-specific ~12-hour genes in BMAL1 knockout mice and BMAL1 knockout mice-specific ~12-hour genes in wild-type mice. (**B, C**) GO analysis of different ~12-hour genes with all mice (**B**) or hepatically expressed (**C**) genes as background.

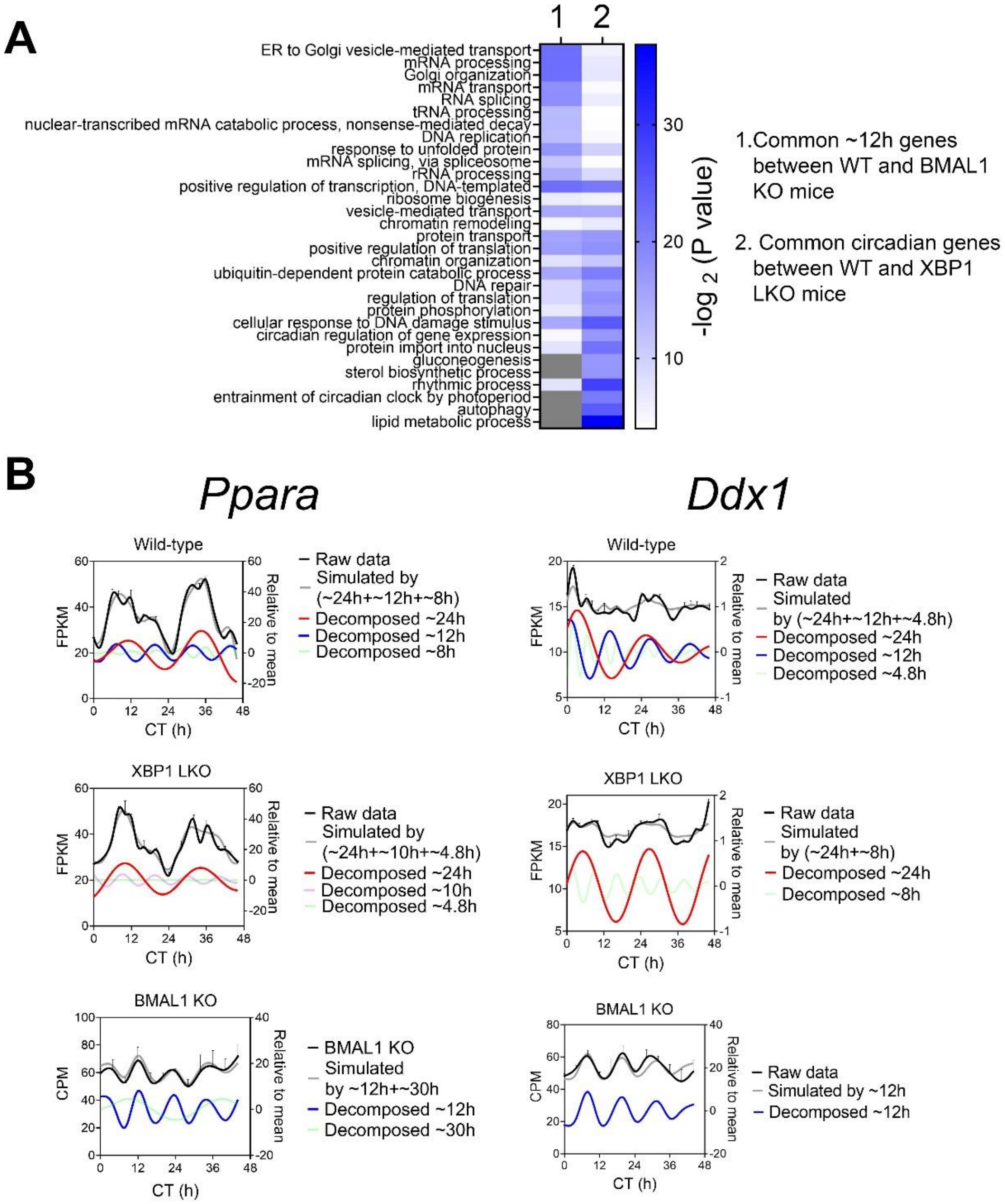


**Figure S4. Many hepatic genes are under dual circadian clock and 12-hour oscillator control.**

(**A**) GO analysis showing enriched biological pathways of common ~12-hour (between wild-type and BMAL1 knockout mice) and circadian (between wild-type and XBP1 LKO mice) using all mice genes as background. (**B**) Eigenvalue/pencil deconvolution of *Ppara* and *Ddx1* temporal gene expression profiles in wild-type, BMAL1 knockout and XBP1 LKO mice. Gray line in each graph illustrates simulated temporal gene expression profile by addition of all superimposed oscillations.

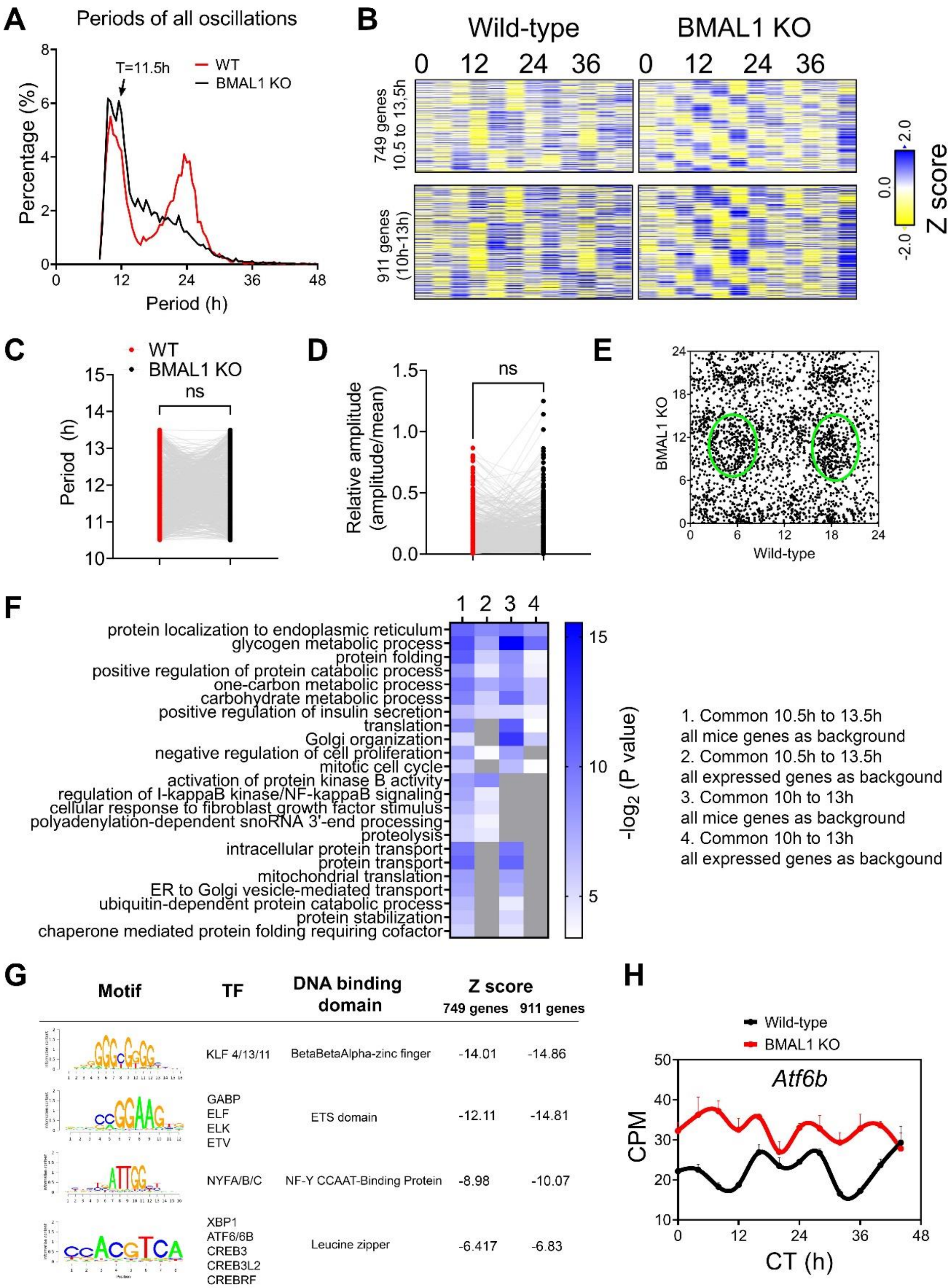
A
Periods of all oscillations
T=11.5h
WT
BMAL1 KO
Percentage (%)
Period (h)
B
Wild-type
BMAL1 KO
0 12 24 36
749 genes 10.5 to 13.5h
911 genes (10h-13h)
Z score
2.0
0.0
-2.0
C
WT
BMAL1 KO
ns
Period (h)
D
ns
Relative amplitude (amplitude/mean)
E
BMAL1 KO
Wild-type
F
1 2 3 4
protein localization to endoplasmic reticulum
glycogen metabolic process
protein folding
positive regulation of protein catabolic process
one-carbon metabolic process
carbohydrate metabolic process
positive regulation of insulin secretion
translation
Golgi organization
negative regulation of cell proliferation
mitotic cell cycle
activation of protein kinase B activity
regulation of I-kappaB kinase/NF-kappaB signaling
cellular response to fibroblast growth factor stimulus
polyadenylation-dependent snoRNA 3'-end processing
proteolysis
intracellular protein transport
protein transport
mitochondrial translation
ER to Golgi vesicle-mediated transport
ubiquitin-dependent protein catabolic process
protein stabilization
chaperone mediated protein folding requiring cofactor
-log2 (P value)
1. Common 10.5h to 13.5h
all mice genes as background
2. Common 10.5h to 13.5h
all expressed genes as backgound
3. Common 10h to 13h
all mice genes as background
4. Common 10h to 13h
all expressed genes as backgound
G
Motif
TF
DNA binding domain
Z score
749 genes
911 genes
KLF 4/13/11
BetaBetaAlpha-zinc finger
-14.01
-14.86
GABP
ELF
ELK
ETV
ETS domain
-12.11
-14.81
NYFA/B/C
NF-Y CCAAT-Binding Protein
-8.98
-10.07
XBP1
ATF6/6B
CREB3
CREB3L2
CREBRF
Leucine zipper
-6.417
-6.83
H
Wild-type
BMAL1 KO
Atf6b
CPM
CT (h)

**Figure S5. Convergence of ~12-hour hepatic transcriptome between wild-type and BMAL1 knockout mice.**

(**A**) Distribution of periods uncovered from all oscillations by the eigenvalue/pencil method from wild-type and BMAL1 knockout mice. (**B**) Heat map of common ~12-hour rhythms of gene expression uncovered by the eigenvalue/pencil method between wild-type and BMAL1 knockout mice. (**C, D**) Period (**C**) and relative amplitude (**D**) of ~12-hour oscillations for 749 genes in wild-type and BMAL1 knockout mice. (**E**) Scatter plot comparing the phases of 749 ~12-hour oscillations in wild-type and BMAL1 knockout mice. (**F**) GO analysis showing enriched biological pathways of common ~12-hour genes using all mice genes or hepatically expressed genes as background. (**G**) Motif analysis of promoter regions of common ~12-hour rhythm genes identified in wild-type and BMAL1 knockout mice. (**H**) Expression of *Atf6b* in wild-type and BMAL1 knockout mice liver.

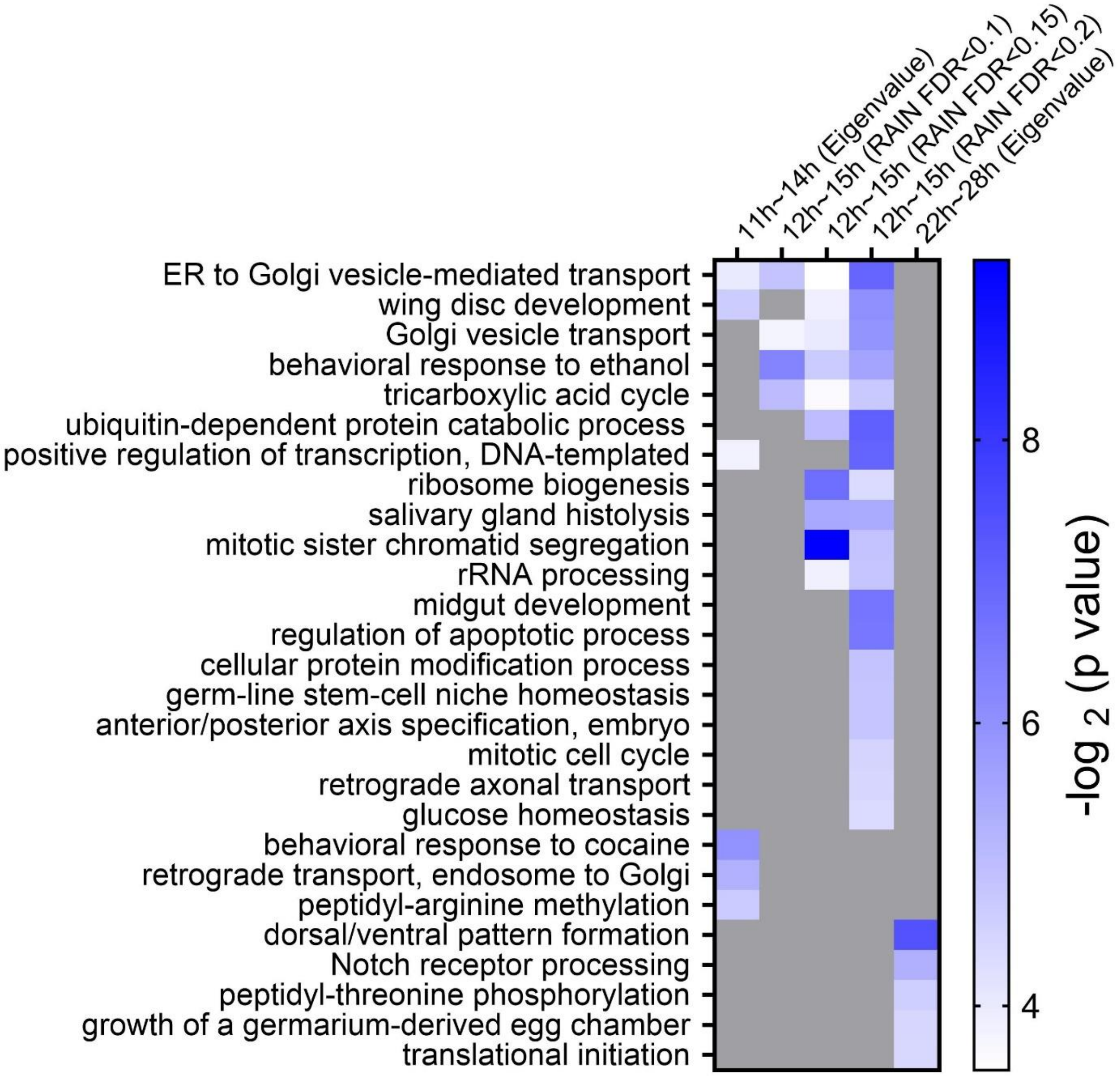


**Figure S6. ~12-hour transcriptome in S2 cells is enriched in protein and RNA metabolism.** GO analysis of ~12-hour and circadian rhythms uncovered by different methods/FDR cut-off in S2 cells using all genes expressed in S2 cells as background.

### Supplemental Tables

**Table S1.** Eigenvalue/pencil deconvolution of temporal gene expression profile of BMAL1 knockout mice.

**Table S2.** RAIN analysis of temporal gene expression profile of BMAL1 knockout mice.

**Table S3.** Eigenvalue/pencil deconvolution of temporal gene expression profile of 141 genes under dual clocks control in wild-type, BMAL1 knockout and XBP1 LKO mice.

**Table S4.** Eigenvalue/pencil deconvolution of temporal gene expression profile of S2 cell.

**Table S5.** RAIN analysis of temporal gene expression profile of S2 cell.
